# Transcriptomic analysis of 40 human and rodent skeletal muscle exerkines

**DOI:** 10.64898/2025.12.19.695001

**Authors:** Hash Brown Taha, Nathan Robbins, Firas Shah-Zoha, Nandhana Vivek, Aleksander Bogoniewski

**Author notes:** Contributed equally to the work (Nathan Robbins, Firas Shah-Zoha). **Author contributions** HBT conceptualized and designed the study. HBT wrote the manuscript. HBT, NR, FZ, NV and AB collected data and checked it for accuracy.

## Abstract

Animal and human studies show that exercise induces organism wide molecular adaptations, many of which are mediated by exerkines which are secreted factors that enable communication between tissues such as skeletal muscle, adipose tissue, liver and the brain. However, the tissue specific responsiveness of individual exerkines and how these responses differ across species, exercise conditions and sexes remain poorly understood. To address this gap, we systematically analyzed skeletal muscle transcriptomic responses of 40 exerkines using three publicly available datasets which include MetaMEx, Extrameta and the MoTrPAC rat training study. We analyzed exerkine specific regulation in humans, mice and rats across acute and chronic exercise as well as inactivity, and determined which responses were conserved, species specific, sex dependent or dependent on exercise duration. Our analysis reveals substantial heterogeneity in skeletal muscle exerkine regulation with only a small subset showing conserved changes across species, while many exerkines exhibited human exclusive, rodent exclusive, acute specific or chronic specific patterns. These results provide a ranked overview of the most exercise responsive skeletal muscle exerkines and highlight the need for multi species and multi condition approaches when selecting exerkines as biomarkers or therapeutic targets.

## Introduction

Animal and human studies strongly support the role of exercise in causing organism-wide improvements in metabolism and health. “Walking is man’s best medicine” is a thousand-year-old quote from Hippocrates [1], illustrating that even with limited understanding of the mechanisms driving various diseases, physical activity and exercise have been recommended as a panacea. Studies have shown that the widespread benefits of exercise can be largely attributed to enhanced intra- and inter-organ communication through molecules termed ‘exerkines’ [2-4]. Indeed, re-infusing exercise-conditioned plasma into non-exercised mice confirms the presence of beneficial bioactive circulatory molecules by offering protection against age-related cognitive decline and enhancing memory [5; 6].

Examples of these exerkines include apelin (APLN), fibroblast growth factor 21 (FGF21), interleukin-6 (IL-6) and brain-derived neurotrophic factor (BDNF) originating from various tissues such as adipose, skeletal muscle, liver and the brain, exerting specific effects in the targeted tissue [2; 3; 7]. However, little is known on the specific molecular response of exerkines within and across different tissues in multiple species (e.g., humans and rodents), and whether there are any intensity-related, time-dependent or sex-specific variations. This is largely attributed to the fact that conducting integrative systems-wide studies requires extensive funding, time and personnel.

Recently, we performed a global multi-omics analysis of 28 exerkines across 20 tissues in exercised 6-month-old rats using data from the Molecular Transducers of Physical Activity Consortium (MoTrPAC), and highlighted several intricate time-dependent, intensity-related and sex-dimorphic responses. Importantly, despite skeletal muscle making up ∼40% of our body weights and thought of as the primary source of exerkine communication in previous studies [2; 8], we discovered that skeletal muscle (gastrocnemius and vastus lateralis) had little exerkine alterations.

To build on these findings and elucidate skeletal muscle-specific exerkine-responsiveness, we gathered a list of 38 known and 2 speculative exerkines and analyzed their skeletal muscle gene expression using two meta-analyses and the MoTrPAC rat 6-month dataset. The two meta-analyses (MetaMEx [9] and Extrameta [10]) pooled gene expression data from published studies to query changes in skeletal muscle transcription responses due to exercise or inactivity, while the MoTrPAC is a global study that uses both human- and rat-based cohorts aiming to uncover the molecular mechanisms by which exercise benefits health [7]. In this study, we only leveraged their publicly available 6month rat data.

Our goal was to pinpoint which skeletal muscle exerkines are most exercise-responsive, elucidate how they vary in multiple species (i.e., humans, mice and rats) and explore whether there are intensity-related, time-dependent and/or sex-dimorphic variations.

## Data extraction and analytical framework

Meta-analytical data analyzed in this study are publicly available from MetaMEx [9] and Extrameta [10], while rat-specific data is publicly available from the MoTrPAC [7]. MetaMEx includes 66 transcriptomic studies for humans and 34 studies for mice, while Extrameta includes 43 studies for humans. Detailed and specific information on each dataset is included in their respective publication. Analyses in this review were made/conducted using https://metamex.serve.scilifelab.se/app/metamex for MetaMEx, https://extrameta.org/ for Extrameta and the MoTrPACRatTraining6moData R package (version 2.0.0) in RStudio (version 4.2.2) for MoTrPAC.

We analyzed a list of 38 known exerkines and 2 speculative exerkines [2; 5; 6; 11-14]. The exerkines include APLN, adiponectin (ADIPOQ), leptin (LEP), meteorin-like (METRNL), growth differentiation factor 15 (GDF15), insulin-like growth factor 1 (IGF-1), FGF21, carboxylesterase 2 and 2C (CES2), PPAR gamma coactivator 1-alpha (PPARGC1A, also known as PGC-1α), fibronectin type III domain-containing protein 5 (FNDC5)/irisin, cathepsin B (CTSB), IL-6, interleukin 7 (IL-7), interleukin 10 (IL-10), interleukin 13 (IL-13), interleukin 15 (IL-15), musclin/osteocrin (OSTN), myostatin (MSTN), follistatin (FST), follistatin like-1 (FSTL1), myonectin (ERFE/CTRP15), decorin (DCN), leukemia inhibitory factor (LIF), secreted protein acidic and rich in cysteine (SPARC), syndecan 4 (SDC4), transforming growth factor beta 1 (TGFβ1), transforming growth factor beta 2 (TGFβ2), angiopoietin 1 (ANGPT1), angiopoietin like-4 (ANGPTL4), fractalkine (CX3CL1), BDNF, neurturin 1 (NTN1), platelet factor 4 (PF4), klotho (KL), fibronectin-1 (FN1), glycosylphosphatidylinositol specific phospholipase D1 (GPLD1), Clusterin (Clu) and fetuin-A (AHSG). We also included the speculative exerkines prosaposin (PSAP) and PSAP like 1 (PSAPL1) given that they are gaining attention as myokines and adipokines [15; 16].

Significance for MetaMEx and Extrameta was determined based on whether the summary point’s error bar crossed the zero line [17]. Timeline-specific significance from MetaMEx was assessed based on a p-value cutoff < 0.05 and whether the logFC was meaningful. For the MoTrPAC, significance was determined using three consecutive conservative methods. P-adjusted values were used to evaluate overall significance. If the P-adjusted value was significant (< 0.05), we used the sex-specific p-values to determine significance. In all cases, if the error bars crossed the zero value, significance was deemed negligible regardless of the p-value [18]. This was we ensure that the specificity of the significance determination for exerkines is high. Acute exercise for MoTrPAC was defined for 1-2 weeks, while chronic exercise was defined as 4-8 weeks.

## Results

Below we provide a brief description and patterns of the analysis. The reader should refer to **Table 1** for more detailed and nuanced comparisons.

**Table 1.**
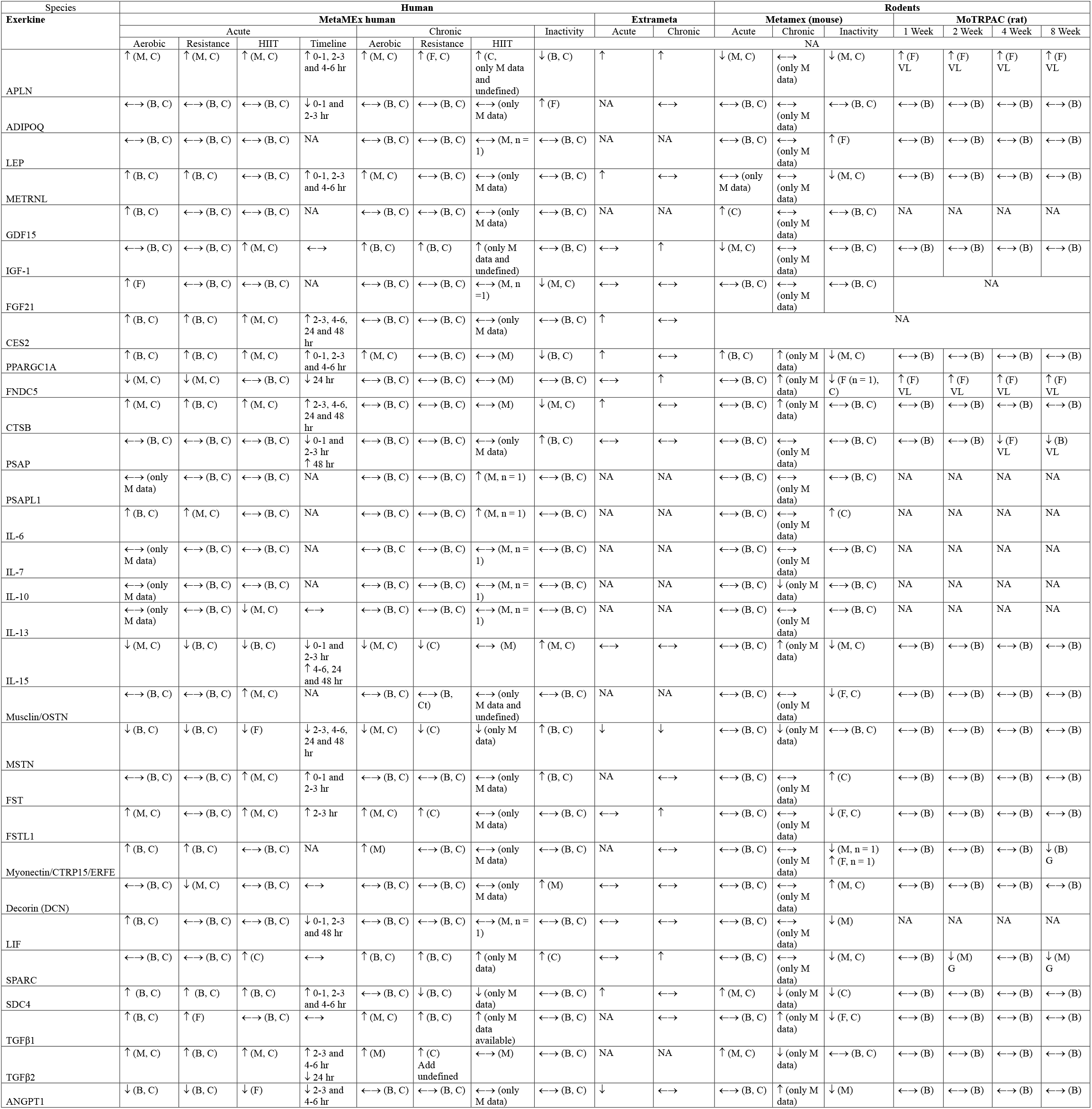
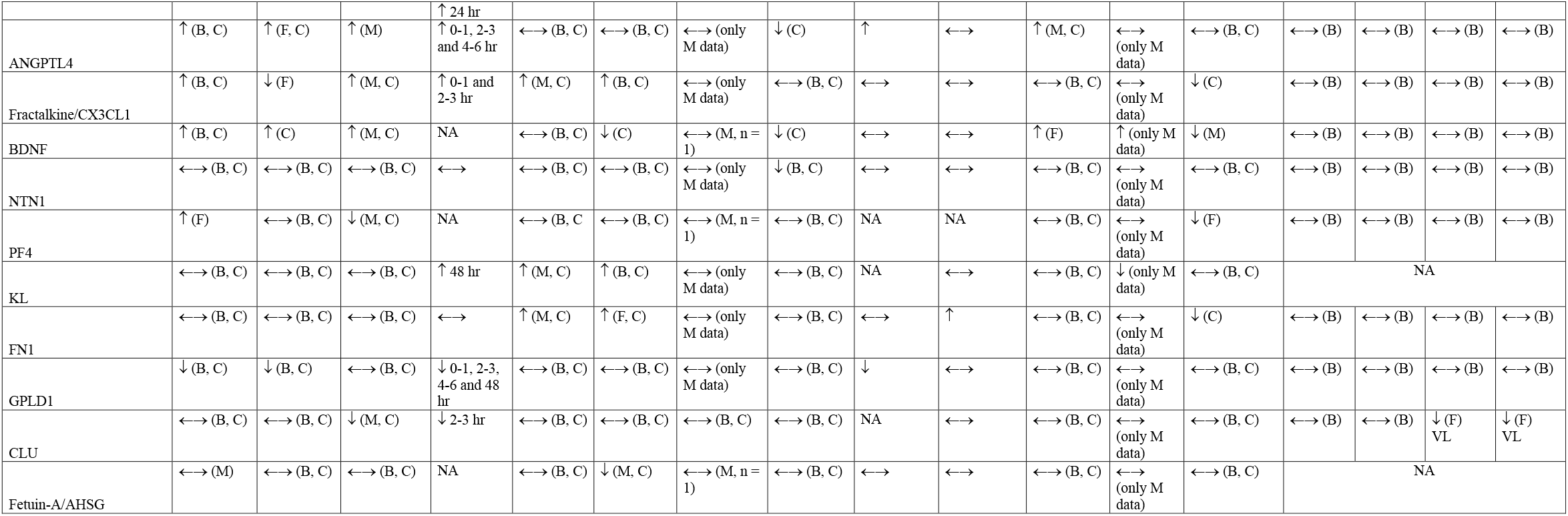
Skeletal muscle-specific exercise responsiveness. MetaMEx data is obtained from https://metamex.serve.scilifelab.se/app/metamex, Extrameta from https://extrameta.org/ and MoTRPAC data from MoTrPACRatTraining6moData R package (version 2.0.0) in RStudio. Exact definition metrics for each condition including aerobic, resistance and high-intensity interval training (HIIT), inactivity, long-term and 1, 2, 4 and 8 weeks of training can be obtained from the respective studies. Note: most studies from MetaMEx and Extrameta included majority male populations. Abbreviations: M – male, F – female, B – both M and F separately, C – combined M and F, VL – vastus latearlis, **APLN** – apelin, **ADIPOQ** – adiponectin, **LEP** – leptin, **METRNL** – meteorin-like, **GDF15** – growth differentiation factor 15, **IGF-1** – insulin-like growth factor 1, **FGF21** – fibroblast growth factor 21, **CES2** – carboxylesterase 2, **PPARGC1A** – **PPAR** gamma coactivator 1-alpha, **FNDC5** – fibronectin type III domain-containing protein 5, **CTSB** – cathepsin B, **PSAP** – prosaposin, **PSAPL1** – prosaposin-like 1, **IL-6** – interleukin 6, **IL-7** – interleukin 7, **IL-10** – interleukin 10, **IL-13** – interleukin 13, **IL-15** – interleukin 15, **Musclin/OSTN** – musclin/osteocrin, **MSTN** – myostatin, **FST** – follistatin, **FSTL** – follistatin-like, **Myonectin/CTRP15/ERFE** – myonectin/C1q TNF related protein 15/erythroferrone, **Decorin** – decorin, **LIF** – leukemia inhibitory factor, **SPARC** – secreted protein acidic and rich in cysteine, **SDC4** – syndecan 4, **TGFβ1** – transforming growth factor beta 1, **TGFβ2** – transforming growth factor beta 2, **ANGPT1** – angiopoietin 1, **ANGPTL4** – angiopoietin-like 4, **Fractalkine/CX3CL1** – fractalkine/C-X3-C motif chemokine ligand 1, **BDNF** – brain-derived neurotrophic factor, **NTN1** – netrin 1, **PF4** – platelet factor 4, **KL** – klotho, **FN1** – fibronectin 1, **GPLD1** – glycosylphosphatidylinositol specific phospholipase D1, **CLU** – clusterin, **Fetuin-A/AHSG** – fetuin-A/alpha-2-HS-glycoprotein.

### Exerkines altered in humans and rodents due to acute exercise

Several exerkines show consistent alterations in response to acute exercise across species. In humans, acute exercise increases APLN, GDF15, IGF-1, PPARGC1A, SDC4, TGFβ2, ANGPTL4, and BDNF. In rodents, findings are more variable: APLN decreases in MetaMEx (primarily in males) but increases in MoTrPAC females at 1–2 weeks; IGF-1 appears to decrease in rodents (MetaMEx only) despite increasing in humans (HIIT exercise only); and GDF15, PPARGC1A, SDC4, TGFβ2, and ANGPTL4 generally increase. BDNF is elevated in humans and rodents (MetaMEx) following acute exercise. These patterns highlight both conserved and species-specific exerkine responses to exercise.

### Exerkines altered in humans and rodents due to chronic exercise

During chronic exercise, only a limited number of skeletal muscles exerkines showed convergent regulation across humans and rodents. APLN and FNDC5 consistently increased in both species, with rodent findings supported by both MetaMEx and MoTrPAC, indicating robust cross species conservation. MSTN and SDC4 also showed concordant decreases in humans and rodents, consistent with its known suppression by training. In contrast, several exerkines displayed opposing regulation between species, including IL15 (decreased in humans, increased in rodents), TGFβ2 (increased in humans, decreased in rodents), BDNF (decreased in humans, increased in rodents), and KL (increased in humans, decreased in rodents). Other molecules such as PPARGC1A, SDC4, and TGFβ1 showed clear regulation in humans but limited or MetaMEx only evidence in rodents, highlighting species specific transcriptional responses. Interestingly, two exerkines showed contrasting results in humans and rodents using MoTrPAC. SPARC and myonectin increased in humans and decreased in rodents. Collectively, aside from APLN and FNDC5, most chronic exercise induced skeletal muscle exerkine changes lacked consistent cross species convergence.

### Exerkines altered in humans and mice due to inactivity

Inactivity leads to a coordinated suppression of several exercise-responsive exerkines across species. Both humans and rodents show decreased APLN, PPARGC1A, and BDNF, alongside increases in FST and DCN. Some molecules display species-specific divergence: IL-15 increases in humans but decreases in rodents, and SPARC increases in humans but decreases in rodents. These patterns highlight shared inactivity-induced molecular signatures as well as important species-dependent regulatory differences.

### Exerkines altered only in humans due to acute exercise

Several exerkines exhibit acute exercise responsiveness uniquely in humans, with no corresponding alterations detected in rodents. Human-specific increases were observed for METRNL, FGF21, CES2, CTSB, IL-6, musclin/OSTN, FST, FSTL1, myonectin, LIF, SPARC, and TGFβ1, while METRNL and SPARC showed partial support from MoTrPAC (e.g., SPARC at 2 weeks). Additional human-specific decreases were noted for FNDC5, IL-13, IL-15, DCN, ANGPT1, GPLD1, and CLU. Some molecules demonstrated mixed or context-dependent effects, including fractalkine and PF4. These findings highlight that several acute exerkine responses may be human-exclusive or substantially more pronounced in humans than in rodents, emphasizing important species differences in early molecular responses to exercise.

### Exerkines altered only in humans due to chronic exercise

A subset of exerkines demonstrated chronic exercise responsiveness exclusively in humans, with no corresponding alterations observed in rodents. Human-only increases included METRNL, IGF-1, IL-6 (notably reported only in a single male cohort), FSTL1, fractalkine, and fibronectin (FN1). Additionally, fetuin-A showed a human-specific decrease. These findings indicate that several chronic exerkine adaptations may be uniquely human or substantially more prominent in humans than in rodents, underscoring important species-specific differences in longer-term molecular responses to exercise.

### Exerkines altered only in rodents due to acute exercise

No exerkines showed acute exercise-specific alterations exclusively in rodents, indicating that all acute rodent responses overlapped with human findings or were not uniquely rodent-specific.

### Exerkines altered only in rodents due to chronic exercise

Chronic exercise produced several rodent-specific exerkine alterations that were not observed in human cohorts. These included a decrease in IL-10, an increase in CTSB (restricted to males), a decrease in PSAP (observed at 4 weeks in females and at 8 weeks in both sexes in MoTrPAC), and an increase in ANGPT1.

### Exerkines altered only in humans due to inactivity

Inactivity produced several human-specific exerkine alterations not observed in rodent models. These included an increase in ADIPOQ (female only), PSAP, and MSTN, and decrease in FGF21, CTSB, ANGPTL4, and NTN1.

### Exerkines altered only in rodents due to inactivity

Rodent-specific analyses revealed a broad suppression of exerkine signaling during inactivity, highlighting a coordinated downregulation across multiple molecular pathways. Several key exerkines consistently decreased, including METRNL, FNDC5, Musclin/OSTN, FSTL1, LIF, SDC4, TGFβ1, ANGPT1, Fractalkine and FN1, indicating reduced myokine, adipokine, and matrix-associated activity in the inactive state. Additional decreases were observed in myonectin with mixed sex-specific responses, and PF4, while IL-6 uniquely increased under inactivity. Notably, leptin increased only in females, suggesting a sex-dependent metabolic response. Overall, these findings show that rodents exhibit a more uniform and widespread suppression of exerkine pathways during inactivity compared with humans, underscoring species-specific differences in how sedentary states modulate molecular signaling.

### Exerkines not altered/low evidence

A small group of exerkines showed no consistent or biologically meaningful changes across exercise or inactivity conditions, including IL-7, IL-10, and PSAPL1, the latter having only a single low-evidence report of increased expression during chronic exercise in MetaMEx.

## Conclusion

In this cross-platform analysis, we demonstrate that skeletal muscle exerkine regulation is highly heterogeneous and strongly dependent on species, exercise duration, and physiological context. Only a very limited subset of exerkines showed true cross species convergence.

APLN and FNDC5 were the most robustly conserved molecules, consistently increasing with both acute and chronic exercise in humans and rodents, supported by evidence from both MetaMEx and MoTrPAC. MSTN also showed concordant suppression across species with chronic training, consistent with its established role as a negative regulator of muscle adaptation. In contrast, the majority of exerkines displayed species divergent or context specific regulation. Several molecules showed opposing responses between humans and rodents, including IL15, TGFβ2, BDNF, KL, SPARC, and myonectin, underscoring fundamental differences in transcriptional adaptation to exercise across species. Notably, most rodent responses outside of APLN and FNDC5 were supported only by MetaMEx, with limited confirmation in MoTrPAC, further emphasizing the need for caution when extrapolating rodent skeletal muscle exerkine data to humans. Acute exercise elicited stronger and more numerous human specific transcriptional responses, whereas rodents showed broader and more uniform suppression of exerkines during inactivity.

Overall, our review demonstrates that muscle-specific exerkine responsiveness cannot be inferred from a single species or exercise condition, and that acute vs chronic exercise activates fundamentally different transcriptional pathways. These results provide a ranked, species-informed reference for the most exercise-sensitive skeletal muscle exerkines and underscore the need for standardized, multi-species, multi-omics validation. In the future, integrating multi-omics approaches [19] across species, tissues, and exercise paradigms will be essential to fully elucidate the complexity of skeletal muscle adaptations and to uncover the full biological and translational potential of skeletal muscle exercise-responsive exerkines.

## Additional information

### Data availability

Data used in the preparation of this article were obtained from MetaMEx, Extrameta and the Molecular Transducers of Physical Activity Consortium (MoTrPAC) MoTrPACRatTraining6moData R package [version 2.0.0].

None.

#### Funding

None.

## Acknowledgements

None.

## Notes

### Competing Interest Statement

The authors have declared no competing interest.

